# A single dose of intravenous human amniotic membrane-derived mesenchymal stem cells limits transmural infarction, reduces fibrosis size, and improves left ventricular systolic function in the myocardial ischemic/reperfusion model of rats

**DOI:** 10.1101/2022.07.30.501939

**Authors:** Jia-Wei Sung, Yen-Ting Yeh, Fu-Shiang Peng, Yi-Chen Chen, Ai-Hsien Li, Shinn-Chih Wu

## Abstract

Despite the advances in coronary reperfusion in acute myocardial infarction (MI), post-MI heart failure is still a large burden of public health. Transmural infarction and extended fibrosis contribute largely to post-MI systolic dysfunction and heart failure, and thus cardioprotective strategies are crucial. Human amniotic membrane-derived mesenchymal stem cells (hAMSCs) have been shown with properties of immunomodulation, anti-inflammation, and low immunogenicity, which make them good candidates for cell therapies. In this study, a myocardial ischemia/reperfusion (I/R) model was established in rats, and hAMSCs were administered via the tail vein during coronary reperfusion. Compared to the control group, the rats receiving hAMSCs during the I/R procedure (hAMSC group) exhibited significantly better left ventricular ejection fractions after MI. Histological examinations of the hearts in hAMSC group showed minimal transmural infarction 4 weeks after the I/R procedure. Compared to the control group, hAMSC group had reduced size of cardiac fibrosis and less thinning of myocardial wall. In conclusion, intravenous hAMSCs limit transmural infarction, reduce fibrosis size, and improve left ventricular systolic function after MI in the animal model.

## Introduction

Myocardial infarction (MI) is a major cause of cardiovascular death around the world (Zaman and Kovoor, 2014). Despite the advances in mechanical reperfusion strategies such as percutaneous coronary intervention (PCI) and coronary artery bypass grafting (CABG), there are still unmet needs of cardioprotection to alleviate myocardial necrosis and ischemia/reperfusion (I/R) injury during MI, which could result in post-MI cardiac remodeling and eventually lead to heart failure. Loss of myocardium and extended fibrosis play critical roles in the pathogenesis of post-MI cardiac remodeling (Hinderer and Schenke-Layland, 2019), and therefore we need cardioprotective strategies in addition to opening the occluded coronary arteries.

Mesenchymal stem cells (MSCs) are multipotent adult stem cells with immunomodulatory and anti-inflammatory properties carrying minimal tumorigenesis potential (Saeedi et al., 2019), which make them good candidates for cell therapies. Recent studies demonstrated that MSCs reduced infarct size, alleviated cardiac fibrosis, and improved post-MI heart function in animal models (Amado et al., 2005; Lee et al., 2009). Therefore, MSC therapy has been one of the potential cardioprotective strategies that seems promising to attenuate LV remodeling after MI. MSCs derive from a variety of different tissues, and human amniotic membrane, which is discarded postpartum as medical waste, is one of good sources of MSCs. Human amniotic membrane-derived mesenchymal stem cells (hAMSCs) are provided with large amounts, easy to harvest, with low immunogenicity, and without ethical issues (Kang et al., 2012; Farhadihosseinabadi et al., 2018; Ragni et al., 2021). Moreover, transplantation of hAMSCs has been demonstrated with encouraging results in prevention of fibrosis and functional improvement in different animal models (Kubo et al., 2015; Cetinkaya et al., 2019). The aim of this study is to investigate the cardioprotective effect of intravenously administered hAMSCs to reduce infarct size and improve cardiac function in the cardiac I/R model of rats.

## Materials and methods

### Isolation and culture of hAMSCs

The samples of amnion membrane were obtained from caesarean sections, and all samples were collected from donors with informed consent. The Ethical Committee of the Far Eastern Memorial Hospital approved the study, which was conducted according to principles of the Helsinki Declaration. Fragments of human amnion membrane were washed in Hanks’ balanced salt solution (HBSS, calcium- and magnesium-free) and digested in 0.05% trypsin-EDTA for 1 hour at 37 °C. Then the membrane was washed with cold HBSS to discard the trypsin digest, transferred to new tubes, and added with digesting solution for 1 hour at 37 °C. The supernatant was collected, added with equal volume of HBSS, and centrifuged at 200 g for 4 minutes.

Harvested cells were cultured in polystyrene culture dishes at 37 °C with an atmosphere of 5 % CO_2_ and in low-glucose DMEM (Gibco Co.) supplemented with 10 % fetal bovine serum (Hyclone Co.), 1% sodium pyruvate (Gibco Co.), 1% penicillin-streptomycin (Gibco Co.), 1 mM NEAA (Gibco Co.), 2 mM GlutaMAX (Gibco Co.), 0.1 mM beta-mercaptoethanol (Gibco Co.), and hEGF 10 ng/mL (Gibco Co.). The cultured medium was refreshed every 3 days. At confluence, the cells were harvested for passage with 1X trypLE (Gibco Co.).

### Animals

Male Sprague Dawley rats at 8 weeks old were purchased from BioLASCO (Taiwan). All experimental protocols were approved by the Institutional Animal Care and Use Committees of National Taiwan University, Taipei, Taiwan.

### The cardiac I/R model of rats

The I/R model was performed as previously described (Chen et al., 2018). In brief, the rats were anesthetized with sodium pentobarbital (80mg/kg, i.p.) and endotracheally ventilated with 2% isoflurane (in pure oxygen). Left-sided thoracotomy was performed by a small incision in the fourth intercostal space, and the left anterior descending coronary artery (LAD) was found and tied for 30 minutes before reperfusion. Ischemia was confirmed by electrocardiography and paleness of myocardium dominated by LAD. The tie on the LAD was released after 30 minutes of ligation to allow reperfusion.

### Echocardiographic measurements

Echocardiography was performed before the I/R procedure and 1, 2, 3, and 4 weeks after the procedure (Figure 4) in anesthetized rats using a PROSPECT ULTRASOUND IMAGING SYSTEM. Left ventricular end-diastolic diameter (LVEDD), left ventricular end-systolic diameter (LVESD), and left ventricular fraction shortening (LVFS) were recorded from parasternal long-axis M-mode images using averaged measurements in 3 to 5 consecutive cardiac cycles in accordance with the American Society of Echocardiography guidelines. Left ventricular end-diastolic and end-systolic volumes (LVEDV and LVESV) were calculated from bidimensional long-axis parasternal views taken through the infarcted area by means of the single-plane area-length method (Moon et al., 2003). Left ventricular ejection fraction (LVEF) was calculated as following: LVEF = (LVEDV-LVESV)/LVEDV) × 100%.

### Pathological assessment

Four weeks after surgery, the rats were euthanized to remove the heart. The heart was simply trimmed to eliminate the atrium after immersion in 10% Neutral buffered formalin (NBF) for 72 hours. The remainder of the heart was cut into four pieces with equal width. These pieces were embedded in paraffin and cut into 5-μm slices, which were stained with H & E staining and Masson trichrome staining. LV anterior wall thickness was measured and averaged. The ratio of fibrotic area to entire LV cross-sectional area was calculated with Image J software (Version 1.51j8, NIH).

### Mesenchymal Trilineage Differentiation Tests of hAMSCs

The hAMSCs were cultured using MesenCult™ Adipogenic Differentiation Kit (STEMCELL TECHNOLOGIES, Inc, Canada) and MesenCult™ Osteogenic Differentiation Kit (STEMCELL TECHNOLOGIES, Inc., Canada) to induce differentiation. Cell pellets (containing 2 × 10^6^) were cultured for 21 days in MesenCult™-ACF Chondrogenic Differentiation Kit (STEMCELL TECHNOLOGIES, Inc, Canada) to induce differentiation. Control cells for all treatments were cultured under normal conditions as previously described. After the induced differentiation stage, the cells were stained with alizarin red to assess osteogenic differentiation, Oil Red O to assess adipogenic differentiation, and Alcian blue for chondrogenic differentiation. Chondrogenic pellets were paraffin-embedded and 3 μm sections were taken, which were then rehydrated and further stained with 1% Alcian blue solution.

### Flow cytometry

hMSCs were identified via flow cytometry with FITC-conjugated antibodies against FITC-anti CD90 (BD Bioscience, Franklin Lakes, NJ, USA, 1:100), FITC-anti CD45 (eBioscience, San Diego, CA, USA, 1:100), FITC-anti CD34 (eBioscience, 1:100), FITC-anti CD11b (eBioscience, 1:100), FITC-anti CD105 (eBioscience, 1:100), FITC-anti CD73 (eBioscience, 1:100), FITC-anti CD14 (eBioscience, 1:100), and FITC-anti mouse IgG as negative control.

### Statistical analysis

All data were expressed as mean ± standard error. In echocardiographic measurement, the paired t-test and unpaired t-test were used for difference comparison. One-way analysis of variance (ANOVA) followed by Scheffé’s post-hoc tests of the least significant difference was used in the comparison of LV anterior wall thickness. P < 0.05 was considered statistically significant.

## Results

### Characterization of hAMSCs

In appearance, hAMSCs displayed properties of MSCs including compact colonies, adhesion, and spindle-shaped morphology (Figure 1). Characterization was done by surface marker analysis using flow cytometry. They were negative for CD34, CD11b, CD45, and HLA-DR and positive for CD105, CD90, CD73, and CD44 mesenchymal markers (Figure 2). They were capable of tri-lineage differentiation into adipocytes, chondrocytes, and osteoblasts in vitro (Figure 3).

**Figure 1.**
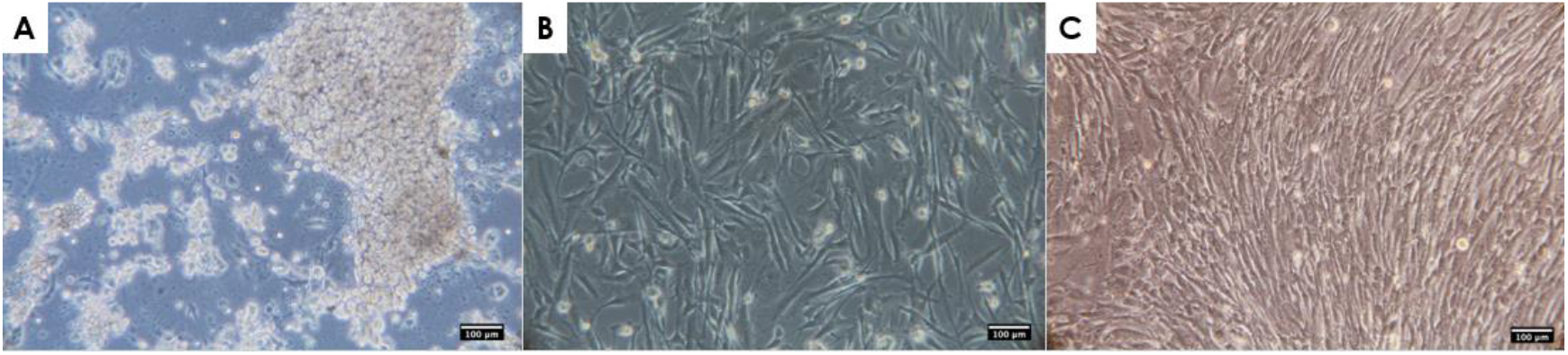
Cell morphology of hAMSCs. (A) hAMSCs on day 1 of primary culture. (B) Fibroblastoid morphology observed at passage 4. (C) Fibroblastoid morphology observed at passage 10. Scale bar = 100 μm.

**Figure 2.**
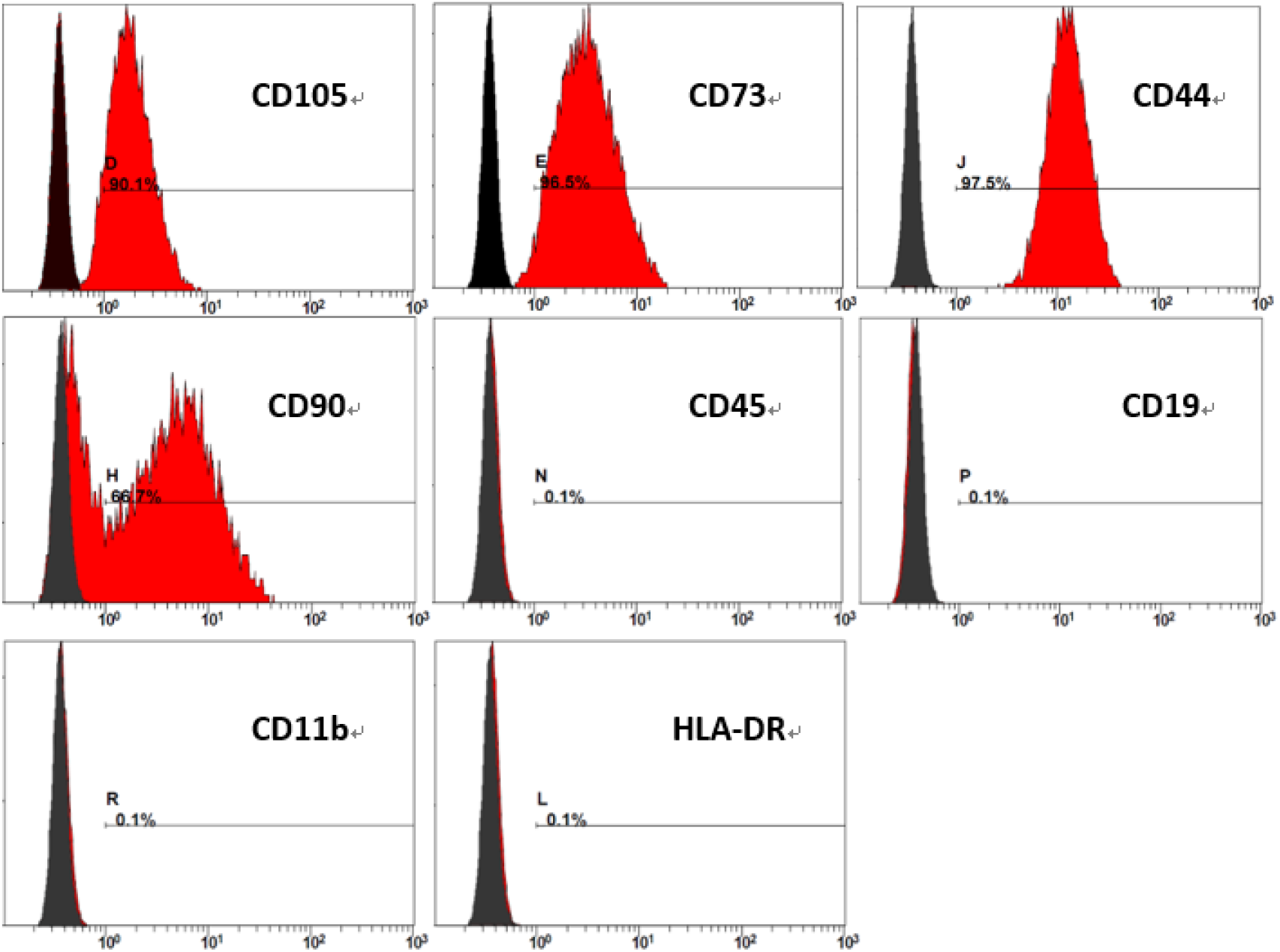
FACS analysis of surface marker expression on hAMSCs at passage 10.

**Figure 3.**
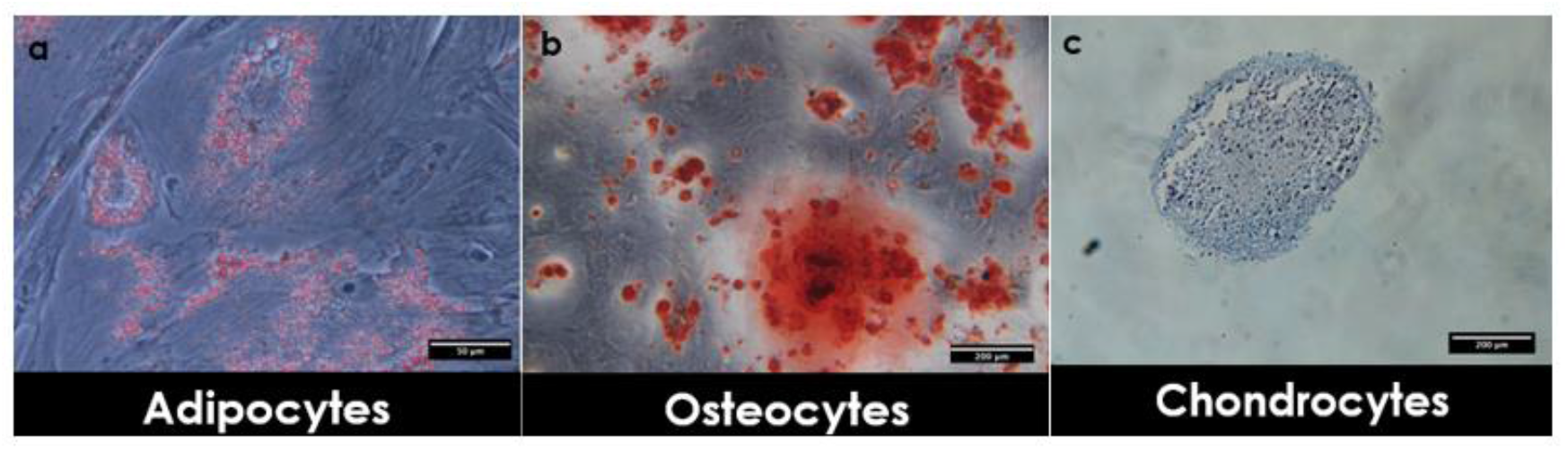
In vitro differentiation of hAMSCs into mesodermal lineages. Cells were cultivated in adipogenic, osteogenic, or chondrogenic differentiation medium. (a) After culture, adipogenic differentiation of cells was shown by intracellular accumulation of neutral lipids stained with Oil Red O solution. Scale bar = 50 μm. (b) Formation of a mineralized matrix after osteogenic differentiation was observed by Alizarin-Red staining. Scale bar = 200 μm. (c) Chondrogenic differentiation was shown by staining with Alcian-Blue. Scale bar = 200 μm.

**Figure 4.**
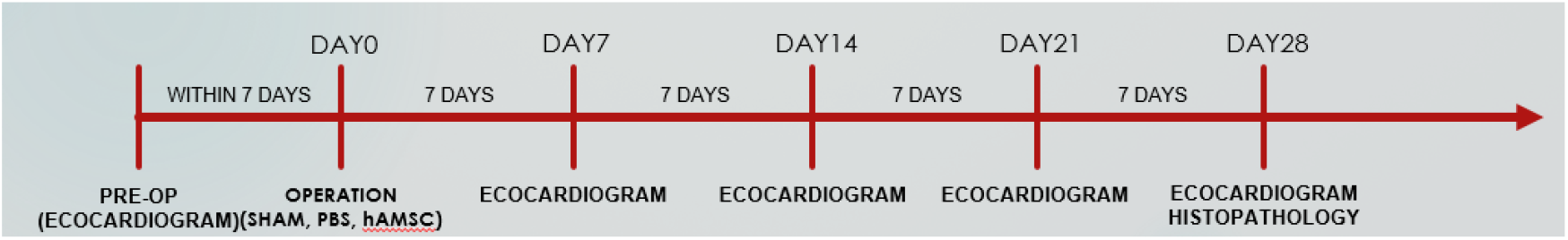
Study flowchart.

### Intravenous administration of hAMSCs improved cardiac function in the I/R model

The rats were randomized into 4 groups. The control group (n=6) was the group with healthy rats that did not undergo any procedure. The sham group (n=6) underwent thoracotomy but the LAD was not ligated. The PBS group (n=6) underwent the whole procedure, and 500 μL PBS without hAMSCs was injected via the tail vein after reperfusion. The hAMSC group (n=6) underwent the same procedure and was injected with 5 × 10^6^ hAMSCs suspended in 500 μL PBS via the tail vein. In 4 weeks after the I/R procedure, no rat died in each group, implying that intravenous treatment of hAMSCs was safe for rats. There was no significant difference in LVEF and LVFS between the sham group and the control group. In the hAMSC group, the LVEF and LVFS were significantly higher than those of the PBS group that received DPBS without hAMSCs throughout the whole 4-week duration (Figure 5 & 6).

**Figure 5.**
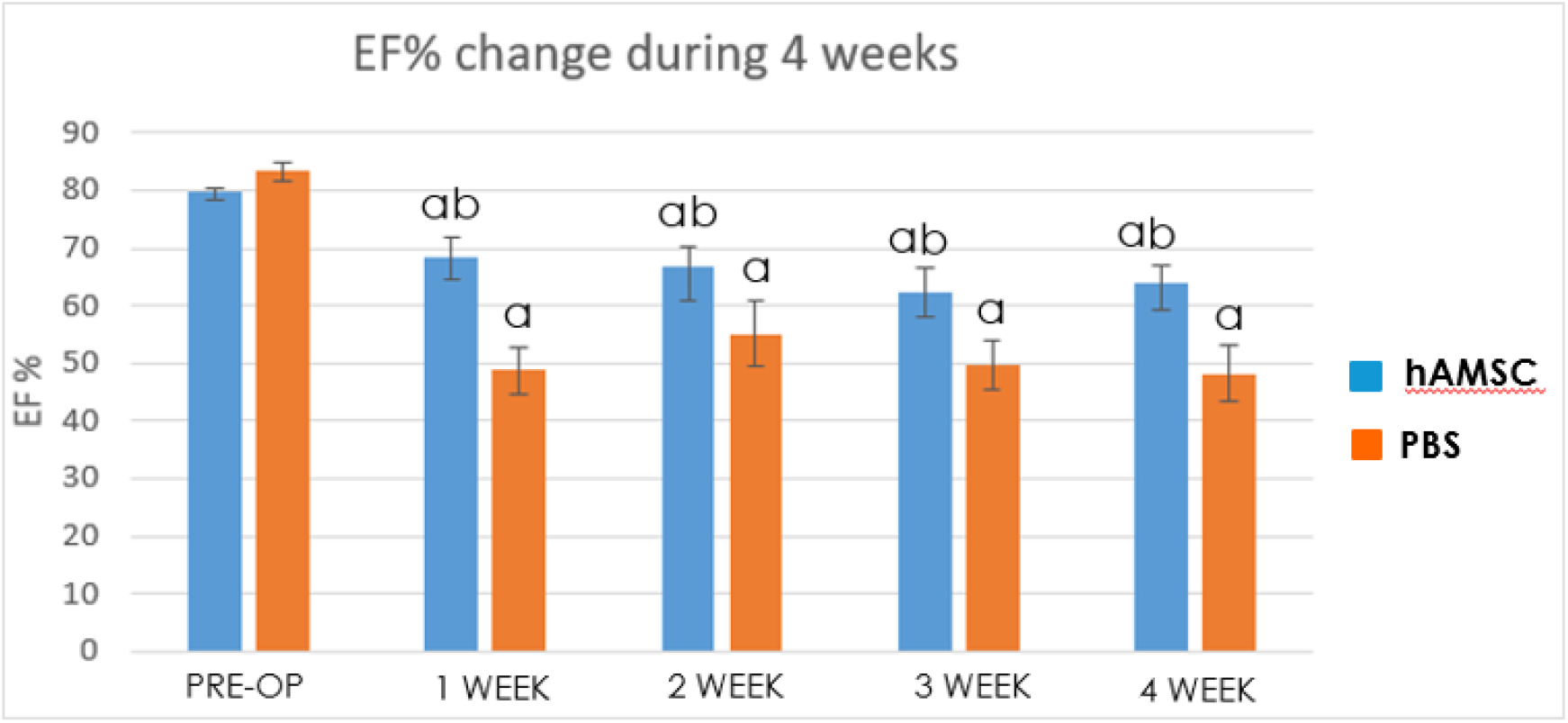
Comparison of LVEF between the hAMSC group and the PBS group during the 4-week period after the procedure. “a” indicates significant difference (P < 0.05) from pre-op, and “b” indicates significant difference (P < 0.05) from the PBS group.

**Figure 6.**
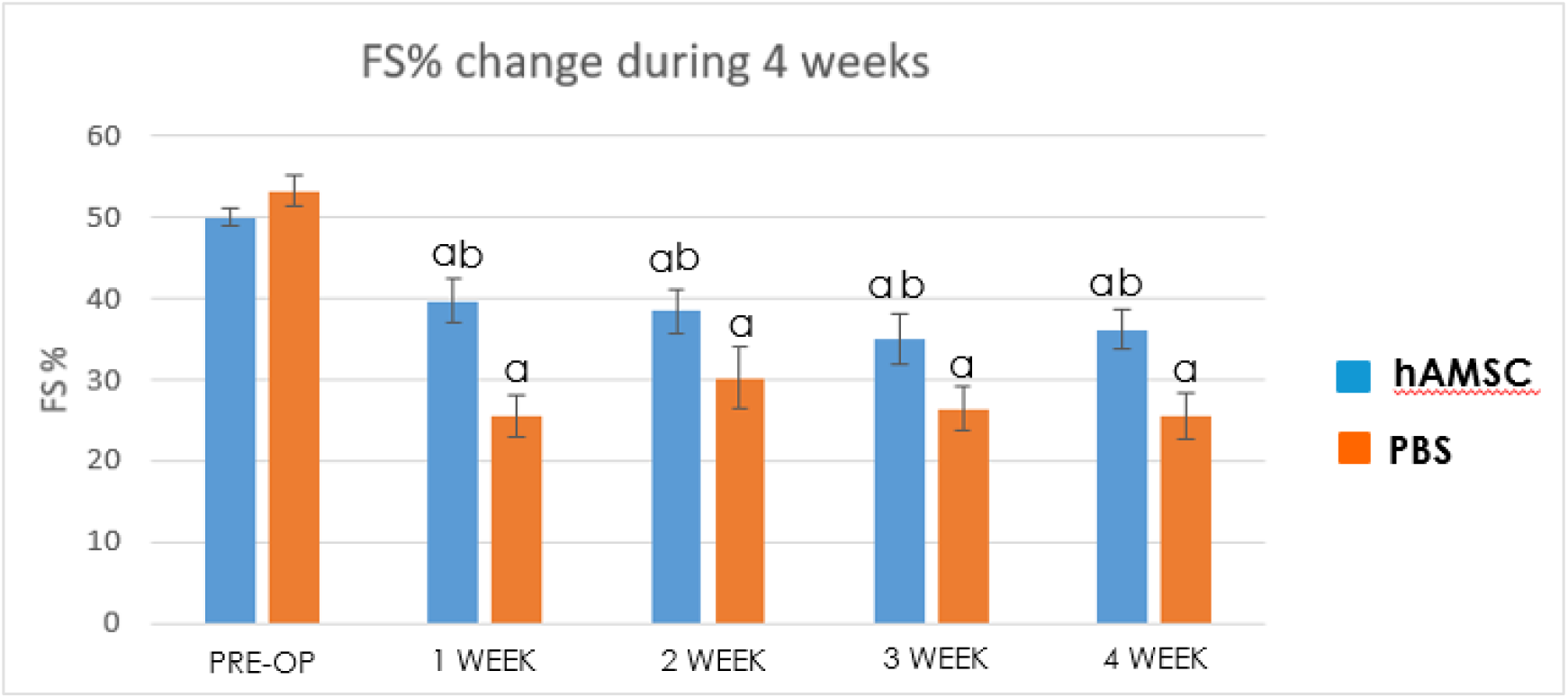
Comparison of LVFS between the hAMSC group and the PBS group during the 4-week period after the procedure. “a” indicates significant difference (P < 0.05) from pre-op. “b” indicates significant difference (P < 0.05) from the PBS group.

### Intravenously administered hAMSCs prevented transmural infarction and reduced LV fibrosis

Cross sections of the harvested hearts were examined with H & E staining and Masson trichrome staining 28 days after the I/R procedure. The overall structure of the LV remained unchanged in the sham group compared to that in the control group. However, an overtly dilated LV and thin LV anterior wall were observed in the PBS group (Figure 7). Quantitative analysis showed that there was no significant difference in LV anterior wall thickness among the control group, the sham group, and the hAMSC group, but the LV anterior wall in the PBS group was significantly thinner than that of the other groups (Figure 8). Fibrotic tissue in LV was marked by blue color with Masson trichrome staining. There was barely blue-stained tissue in the control group and the sham group. In the PBS group, extensive transmural fibrosis of LV wall could be observed. Notably, there was very little transmural fibrosis in the hAMSC group, and most of the fibrotic tissue was limited in subendocardial regions (Figure 9). Quantitative analysis showed the ratio of fibrotic area to total cross sectional area of LV was significantly reduced in the hAMSC group compared to that in the PBS group (Figure 10).

**Figure 7.**
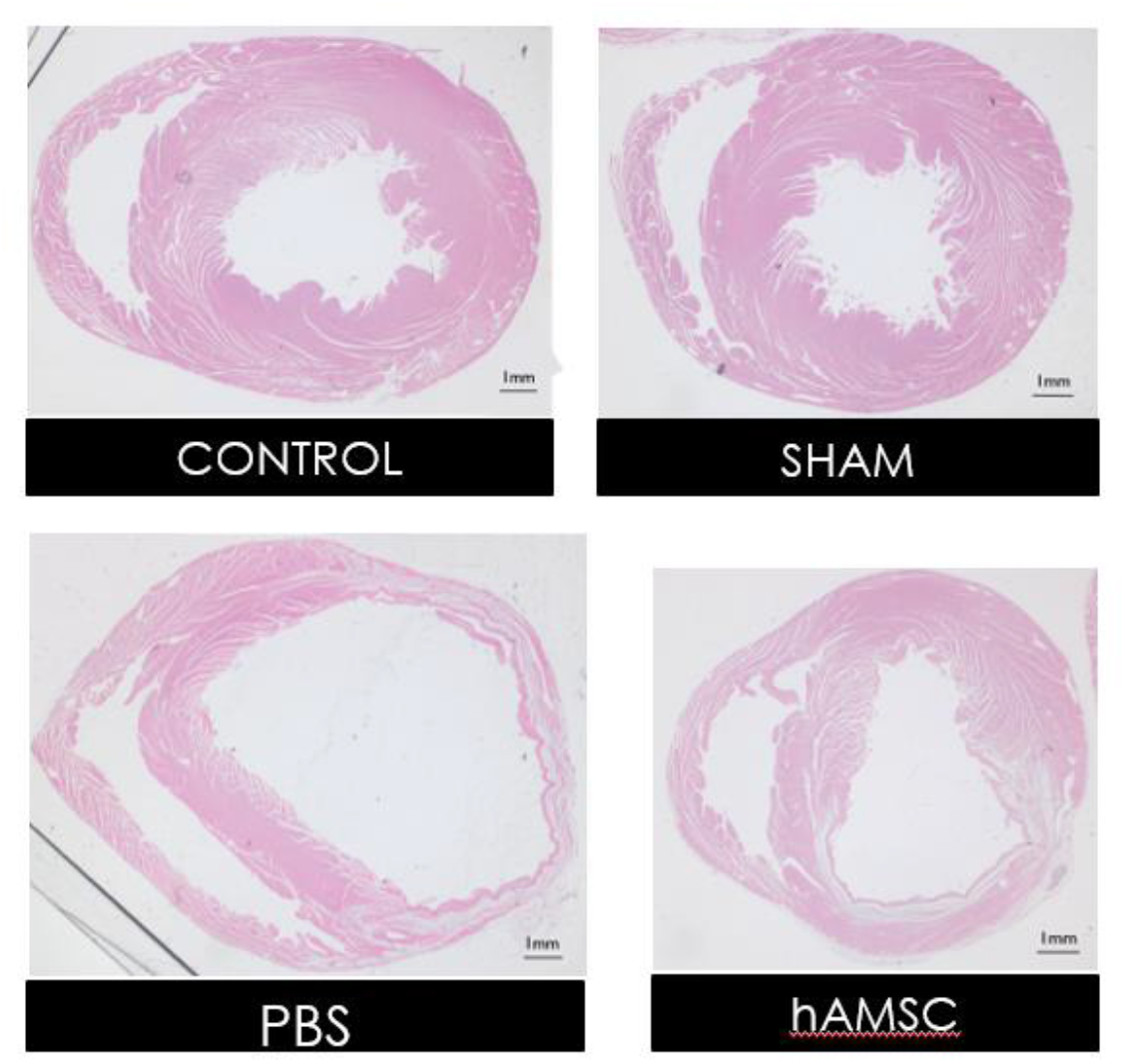
Representative LV sections of the harvested hearts at midventricular level with H & E staining. Scale bar = 1 mm.

**Figure 8.**
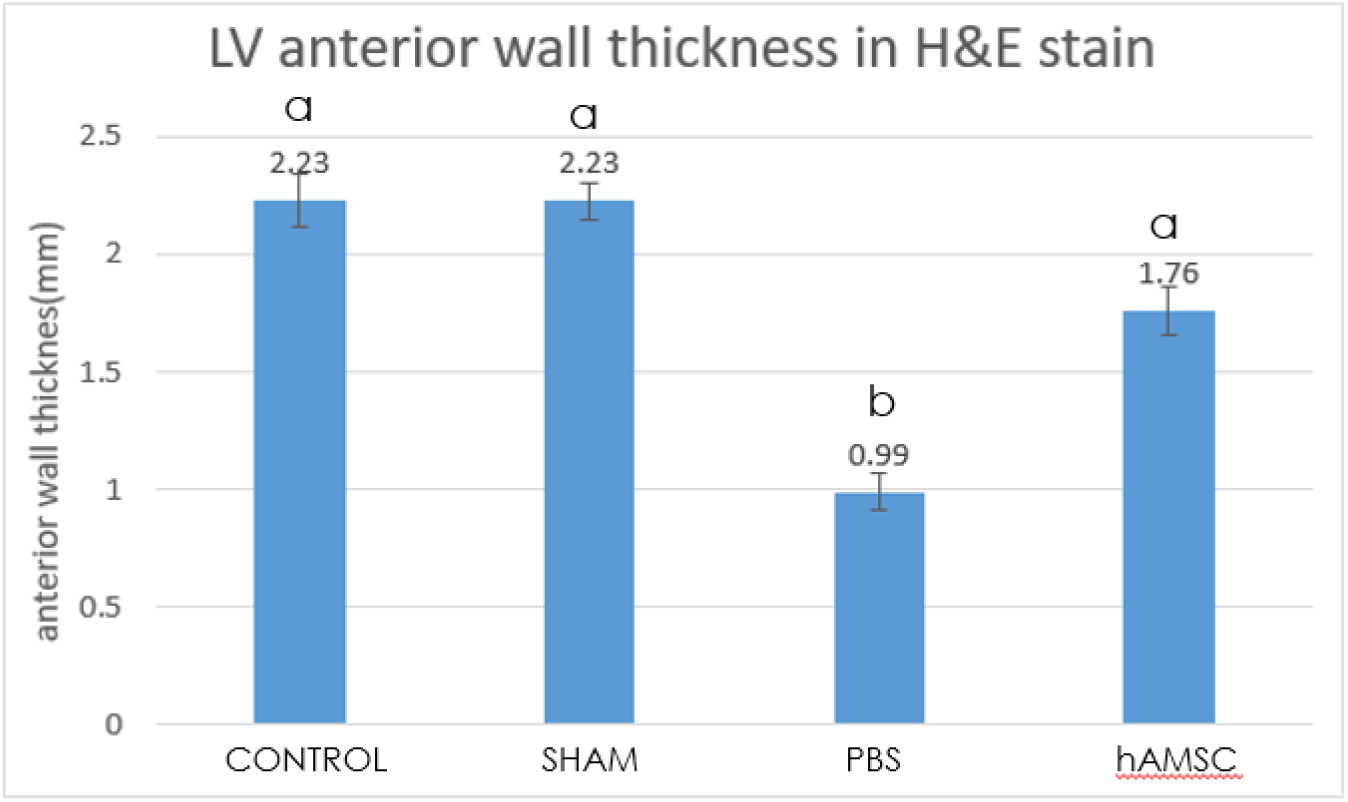
Averaged LV anterior wall thickness of the harvested hearts. “a” indicates significant difference (P < 0.05) from the PBS group. “b” indicates significant difference (P < 0.05) from the control group.

**Figure 9.**
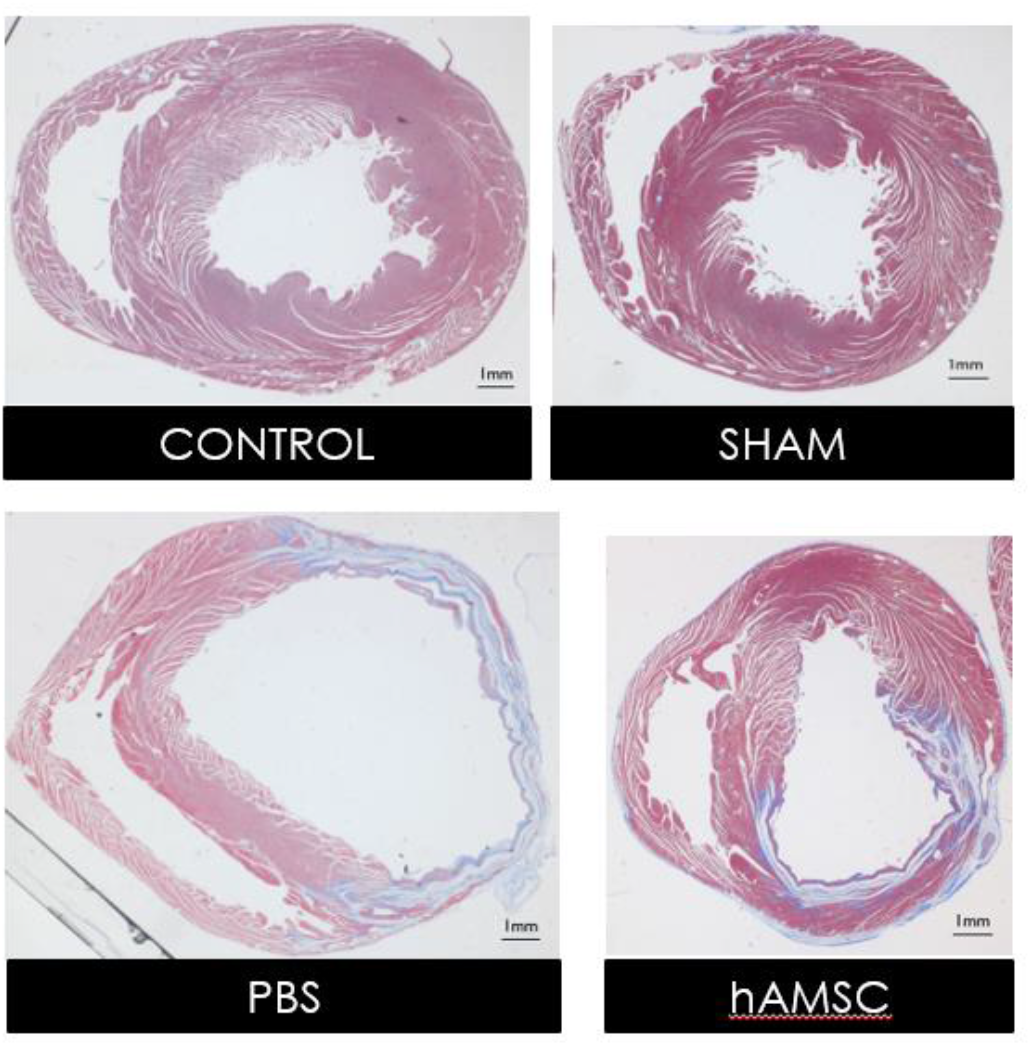
Representative LV sections of the harvested hearts at midventricular level with Masson trichrome staining. Scale bar = 1 mm.

**Figure 10.**
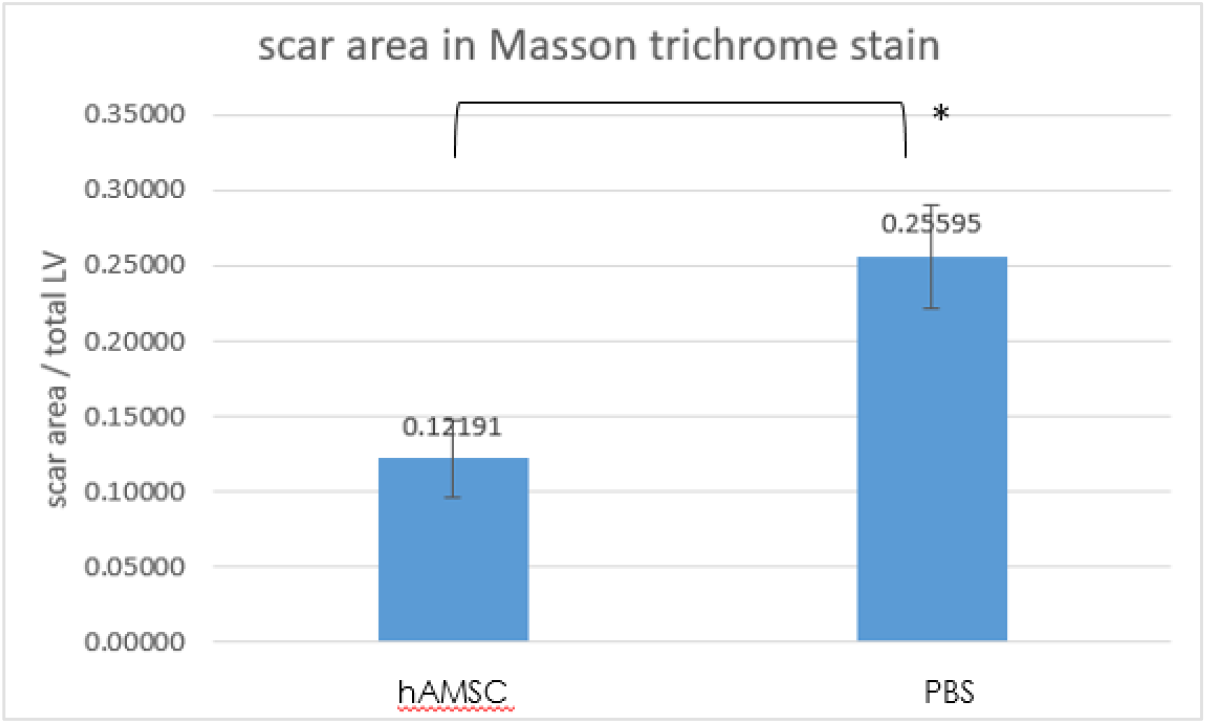
The ratio of fibrotic area to total cross sectional area of LV was significantly reduced in hAMSC group. * P < 0.05

## Discussion

In clinical practice, LV transmural infarction is commonly seen when a coronary artery is totally obstructed in the setting of acute myocardial infarction (AMI) even if the culprit vessel is timely opened with primary PCI. Unfortunately, transmural infarction often leads to consequences including infarct expansion, excessive fibrosis, LV aneurysms, and eventually heart failure (Jenca et al., 2021). In this study, we successfully established a rat I/R model that mimics the clinical setting of AMI undergoing timely coronary reperfusion. In the PBS group, extensive LV transmural fibrosis and marked loss of LV myocardium developed despite rapid restoration of blood flow of the ligated LAD in the I/R procedure. On the other hand, histological examination in the hAMSC group showed very little transmural fibrosis, and the LV myocardium was largely preserved since there was no significant thinning of the anterior wall. The data of serial echocardiographic measurement were consistent with histological findings. The hAMSC group had significantly better LV systolic function compared to the PBS group. These results demonstrated a single dose of intravenous hAMSCs at the moment of coronary reperfusion could prevent transmural infarction and alleviate LV remodeling, which translated into better post-MI LV systolic function.

Amniotic membrane (AM), traditionally a medical waste, has been employed as scaffolding material in tissue engineering applications such as skin dressing or the graft for corneal treatment (Meller et al., 2011; Salehi et al., 2015), which have shown the safety of allogeneic use of AM tissue. In recent years, hAMSCs, isolated from human AM, have been demonstrated with good differentiation potentials, immunomodulatory properties, and low immunogenicity (Farhadihosseinabadi et al., 2018; Kang et al., 2012; Ragni et al., 2021). Studies demonstrated that xenotransplantation of hAMSCs resulted in minimal acute graft rejection reaction, making them excellent candidates of cell therapy (Qiang et al., 2016; Cui et al., 2018; Hua et al., 2019; Yin et al., 2019).

Clinical studies have shown that autologous bone marrow cells or MSCs exhibited some benefits in AMI patients (Martin-Rendon et al., 2008; Narita and Suzuki, 2015; Kim et al., 2018). However, the therapeutic effect of autologous cell therapy could be largely limited by several factors. The timing of cell administration in clinical studies was likely suboptimal since procedures of harvest, culture, and expansion of autologous cells take at least several days, which might have already passed the “golden time” of cardioprotection. The dosage of administered cells was limited to the donor factor and short expansion time, which also made quality control barely possible. Allogeneic hAMSCs could be the solution of above problems since their property of low immunogenicity and proven safety of allogeneic use of human AM. Given the abundant sources of hAMSCs, which could be collected from daily caesarean section procedures and made into a prestored product, the dosage and quality of cells would be problems no more. Furthermore, intracoronary or intravenous hAMSCs could be given right at the time of coronary reperfusion to achieve best cardioprotective effect, which was demonstrated in this animal study to prevent transmural infarction, alleviate myocardial loss, improve LV systolic function, and possibly reduce heart failure. Future studies are warranted.

## Conclusion

A single dose of intravenous hAMSCs administered at the time of coronary reperfusion prevented transmural infarction, alleviated myocardial loss, and improved LV systolic function in the rat I/R model.

## Supporting information

cover letter

## References

Amado, L. C., A. P. Saliaris, K. H. Schuleri, M. St John, J. S. Xie, S. Cattaneo, D. J. Durand, T. Fitton, J. Q. Kuang, G. Stewart, S. Lehrke, W. W. Baumgartner, B. J. Martin, A. W. Heldman, and J. M. Hare. (2005). Cardiac repair with intramyocardial injection of allogeneic mesenchymal stem cells after myocardial infarction. Proc. Natl. Acad. Sci. U S A 102:11474–11479. doi: 10.1073/pnas.0504388102

Cetinkaya, B., G. Unek, D. Kipmen-Korgun, S. Koksoy, and E. T. Korgun. (2019). Effects of Human Placental Amnion Derived Mesenchymal Stem Cells on Proliferation and Apoptosis Mechanisms in Chronic Kidney Disease in the Rat. Int. J. Stem Cells 12:151–161. doi: 10.15283/ijsc18067

Chen, J., D. K. Ceholski, I. C. Turnbull, L. Liang, and R. J. Hajjar. (2018). Ischemic Model of Heart Failure in Rats and Mice. Methods Mol. Biol. 1816:175–182. doi: 10.1007/978-1-4939-8597-5_13

Cui, P., H. Xin, Y. Yao, S. Xiao, F. Zhu, Z. Gong, Z. Tang, Q. Zhan, W. Qin, Y. Lai, X. Li, Y. Tong, and Z. Xia. (2018). Human amnion-derived mesenchymal stem cells alleviate lung injury induced by white smoke inhalation in rats. Stem Cell Res. Ther. 9:101. doi: 10.1186/s13287-018-0856-7

Farhadihosseinabadi, B., M. Farahani, T. Tayebi, A. Jafari, F. Biniazan, K. Modaresifar, H. Moravvej, S. Bahrami, H. Redl, L. Tayebi, and H. Niknejad. (2018). Amniotic membrane and its epithelial and mesenchymal stem cells as an appropriate source for skin tissue engineering and regenerative medicine. Artif. Cells Nanomed. Biotechnol. 46:431–440. doi: 10.1080/21691401.2018.1458730

Hinderer, S., and K. Schenke-Layland. (2019). Cardiac fibrosis - A short review of causes and therapeutic strategies. Adv. Drug Deliv. Rev. 146:77–82. doi: 10.1016/j.addr.2019.05.011

Hua, D., Z. Ju, X. Gan, Q. Wang, C. Luo, J. Gu, and Y. Yu. (2019). Human amniotic mesenchymal stromal cells alleviate acute liver injury by inhibiting the proinflammatory response of liver resident macrophage through autophagy. Ann. Transl. Med. 7:392. doi: 10.21037/atm.2019.08.83

Jenca, D., V. Melenovsky, J. Stehlik, V. Stanek, J. Kettner, J. Kautzner, V. Adamkova, and P. Wohlfahrt. (2021). Heart failure after myocardial infarction: incidence and predictors. ESC Heart Fail. 8:222–237. doi: 10.1002/ehf2.13144

Kang, J. W., H. C. Koo, S. Y. Hwang, S. K. Kang, J. C. Ra, M. H. Lee, and Y. H. Park. (2012). Immunomodulatory effects of human amniotic membrane-derived mesenchymal stem cells. J. Vet. Sci. 13:23–31. doi: 10.4142/jvs.2012.13.1.23

Kim, S. H., J. H. Cho, Y. H. Lee, J. H. Lee, S. S. Kim, M. Y. Kim, M. G. Lee, W. Y. Kang, K. S. Lee, Y. K. Ahn, M. H. Jeong, and H. S. Kim. (2018). Improvement in Left Ventricular Function with Intracoronary Mesenchymal Stem Cell Therapy in a Patient with Anterior Wall ST-Segment Elevation Myocardial Infarction. Cardiovasc. Drugs Ther. 32:329–338. doi: 10.1007/s10557-018-6804-z

Kubo, K., S. Ohnishi, H. Hosono, M. Fukai, A. Kameya, R. Higashi, T. Yamada, R. Onishi, K. Yamahara, H. Takeda, and N. Sakamoto. (2015). Human Amnion-Derived Mesenchymal Stem Cell Transplantation Ameliorates Liver Fibrosis in Rats. Transplant. Direct 1:e16. doi: 10.1097/TXD.0000000000000525

Lee, R. H., A. A. Pulin, M. J. Seo, D. J. Kota, J. Ylostalo, B. L. Larson, L. Semprun-Prieto, P. Delafontaine, and D. J. Prockop. (2009). Intravenous hMSCs improve myocardial infarction in mice because cells embolized in lung are activated to secrete the anti-inflammatory protein TSG-6. Cell Stem Cell 5:54–63. doi: 10.1016/j.stem.2009.05.003

Martin-Rendon, E., S. J. Brunskill, C. J. Hyde, S. J. Stanworth, A. Mathur, and S. M. Watt. (2008). Autologous bone marrow stem cells to treat acute myocardial infarction: a systematic review. European Heart Journal 29:1807–1818. doi: 10.1093/eurheartj/ehn220

Meller, D., M. Pauklin, H. Thomasen, H. Westekemper, and K. P. Steuhl. (2011). Amniotic membrane transplantation in the human eye. Dtsch. Arztebl. Int. 108:243–248. doi: 10.3238/arztebl.2011.0243

Moon, C., M. Krawczyk, D. Ahn, I. Ahmet, D. Paik, E. G. Lakatta, and M. I. Talan. (2003). Erythropoietin reduces myocardial infarction and left ventricular functional decline after coronary artery ligation in rats. Proc. Natl. Acad. Sci. U S A 100:11612–11617. doi: 10.1073/pnas.1930406100

Narita, T., and K. Suzuki. (2015). Bone marrow-derived mesenchymal stem cells for the treatment of heart failure. Heart Fail. Rev. 20:53–68. doi: 10.1007/s10741-014-9435-x

Qiang, Y., G. Liang, and L. Yu. (2016). Human amniotic mesenchymal stem cells alleviate lung injury induced by ischemia and reperfusion after cardiopulmonary bypass in dogs. Lab. Invest. 96:537–546. doi: 10.1038/labinvest.2016.37

Ragni, E., A. Papait, C. Perucca Orfei, A. R. Silini, A. Colombini, M. Vigano, F. Libonati, O. Parolini, and L. de Girolamo. (2021). Amniotic membrane-mesenchymal stromal cells secreted factors and extracellular vesicle-miRNAs: Anti-inflammatory and regenerative features for musculoskeletal tissues. Stem Cells Transl. Med. 10:1044–1062. doi: 10.1002/sctm.20-0390

Saeedi, P., R. Halabian, and A. A. Imani Fooladi. (2019). A revealing review of mesenchymal stem cells therapy, clinical perspectives and Modification strategies. Stem Cell Investig. 6:34. doi: 10.21037/sci.2019.08.11

Salehi, S. H., K. As’adi, S. J. Mousavi, and S. Shoar. (2015). Evaluation of Amniotic Membrane Effectiveness in Skin Graft Donor Site Dressing in Burn Patients. Indian J. Surg. 77:427–431. doi: 10.1007/s12262-013-0864-x

Yin, L., Z. X. Zhou, M. Shen, N. Chen, F. Jiang, and S. L. Wang. (2019). The Human Amniotic Mesenchymal Stem Cells (hAMSCs) Improve the Implant Osseointegration and Bone Regeneration in Maxillary Sinus Floor Elevation in Rabbits. Stem Cells Int. 2019:9845497. doi: 10.1155/2019/9845497

Zaman, S., and P. Kovoor. (2014). Sudden cardiac death early after myocardial infarction: pathogenesis, risk stratification, and primary prevention. Circulation 129:2426–2435. doi: 10.1161/CIRCULATIONAHA.113.007497

