## Supplementary material for "A single dose of intravenous human amniotic membrane-derived mesenchymal stem cells limits transmural infarction, reduces fibrosis size, and improves left ventricular systolic function in the myocardial ischemic/reperfusion model of rats": cover letter

Dear Editor in Chief,

We would like to submit the enclosed manuscript entitled “A single dose of intravenous human amniotic membrane-derived mesenchymal stem cells limits transmural infarction, reduces fibrosis size, and improves left ventricular systolic function in the myocardial ischemic/reperfusion model of rats”, which we wish to be considered for publication in Disease Models & Mechanisms. There is no conflict of interests in this manuscript. I would like to declare on behalf of my co-authors that the work described was original research that has not been published previously, and not under consideration for publication elsewhere, in whole or in part. All the authors listed have approved the manuscript that is enclosed.
In this study, we investigated the cardioprotective effect of intravenous human amnion-derived mesenchymal stem cells (hAMSCs) in the myocardial ischemia/reperfusion (IR) model of rats. The results demonstrated that a single dose of intravenous hAMSCs limited transmural infarction, reduced infarct size, and improved left ventricular systolic function, which is crucial in prevention of heart failure. I hope this manuscript is suitable for the upcoming special issue “Moving Heart Failure to Heart Success: Mechanisms, Regeneration & Therapy”.
We deeply appreciate your consideration of our manuscript and look forward to hearing from you. If you have any queries, please don’t hesitate to contact me at the address below.

Yours sincerely,
Shinn-Chih Wu
